# The Homeostatic Logic of Reward

**DOI:** 10.1101/242974

**Authors:** Tobias Morville, Karl Friston, Denis Burdakov, Hartwig R. Siebner, Oliver J. Hulme

## Abstract

Energy homeostasis depends on behavior to predictively regulate metabolic states within narrow bounds. Here we review three theories of homeostatic control and ask how they provide insight into the circuitry underlying energy homeostasis. We offer two contributions. First, we detail how control theory and reinforcement learning are applied to homeostatic control. We show how these schemes rest on implausible assumptions; either via circular definitions, unprincipled drive functions, or by ignoring environmental volatility. We argue active inference can elude these shortcomings while retaining important features of each model. Second, we review the neural basis of energetic control. We focus on a subset of arcuate subpopulations that project directly to, and are thus in a privileged position to opponently modulate, dopaminergic cells as a function of energetic predictions over a spectrum of time horizons. We discuss how this can be interpreted under these theories, and how this can resolve paradoxes that have arisen. We propose this circuit constitutes a homeostatic-reward interface that underwrites the conjoint optimisation of physiological and behavioural homeostasis.

## The problem of homeostatic control

A remarkable feature of physiological systems is their stability. Most physiological variables are regulated within narrow bounds by operational and computational processes collectively known as homeostasis (Cannon 1932). The mechanistic complexity of homeostasis extends beyond simple negative feedback control and embodies a wide spectrum of hierarchically organised physiological control structures, molecule to agent, operating over a multitude of timescales, milliseconds to months (Carpenter 2004). Homeostatic control is often framed as the regulation of variables around a fixed set point, the achievement of which upholds a physiological equilibrium (Cannon 1932). Fixed set points, however useful they are as abstractions, are biologically implausible. Indeed, allostasis (under some definitions, e.g. Sterling 2012, Stephan et al. 2016) refers to the dynamic process by which homeostatic equilibria shift. For instance moving set points could occur through the transient modulations of stress, digestion, or arousal (Peters et al 2017), through to longer timescales of circadian or circannual rhythms, developmental or reproductive phases (this form of predictive regulation is also known as *rheostasis* and a great many other names, see Woods & Ramsay 2007). However, since it has been argued that predictive control does not distinguish between allostasis and homeostasis (Woods & Ramsay 2007), we use the term homeostasis in this broadest sense, to encompass predictive control. In other words, homoeostasis here subsumes classical homeostatic reflexes and hierarchically embellished allostatic control. Under this nomenclature, the raison d’être of homeostasis is not stability per se, but rather dynamically adjusting internal states to fall within the ranges that afford organismal survival (Fig 1b; Sterling 2012).

**Figure 1.**
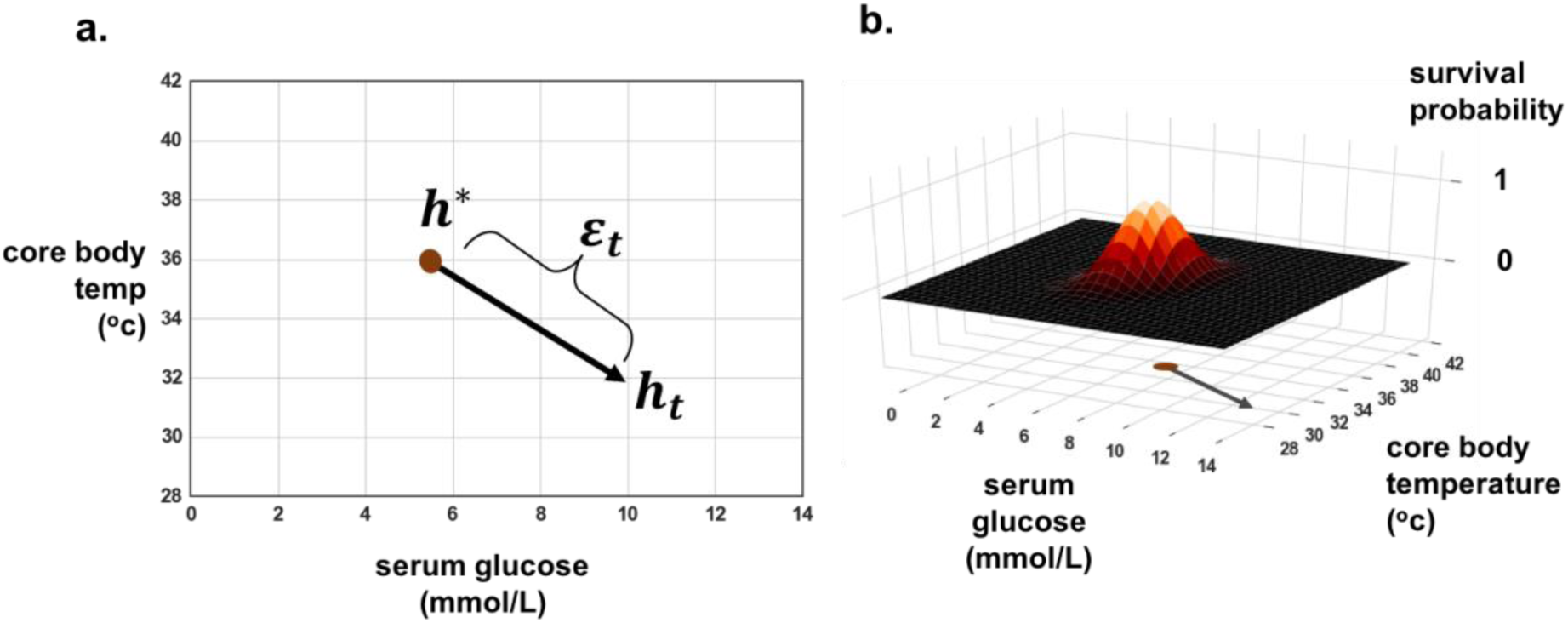
Homeostatic state space and survival probability. **a**, Schematic showing a simple 2-dimensional homeostatic state space, where *h\** denotes a set point, *h_t_* current state at time *t*, and the error between them defined here as the absolute Euclidean distance ε*_t_*,. **b**, Shows a survival probability surface, depicted over the same homeostatic state space, thus highlighting the relation between homeostatic error and the conditional probability of survival (over some time interval), given the occupation of that homeostatic state.

For all motile agents, effective homeostatic control results from the interplay among automated physiological processes (henceforth referred to as physiological homeostasis) and overt behaviour (henceforth, behavioural homeostasis). The coordinated mechanisms of physiological homeostasis are insufficient to perpetuate survival. In an indolent (inactive or sessile) organism, the incessant activity of basal metabolic processes results in the continuous drift of vital homeostatic variables, as time passes. These excursions cannot be mitigated by the coordinated mechanisms of physiological homeostasis alone. The homeostatic error, defined as the distance of the current homeostatic state (physiological state) from any set point (Fig. 1a) can only redressed by behavioural exchange with the external environment (hunting, seeking warmth, micturition etc.). Thus, homeostatic control consists of tracking, estimating, and predicting homeostatic errors, and simultaneously prioritizing and generating the appropriate physiological and behavioural responses to minimize those errors.

In the terms defined above, this entails a conjoint optimisation of both physiological and behavioural homeostasis. From a computational perspective, this is a challenging problem for many reasons: All natural habitats are complex, uncertain, labile, and often precarious. The resources of utility for reducing homeostatic error are typically sparsely distributed in time and space. Internal states have to be inferred accurately, or at least as accurately as their survival hazards mandate. Each internal state has its own dynamics and uncertainties, so any control mechanism has to contend with variables interacting over multiple timescales, often with different degrees of synergy, antagonism, and over different scales of lag. Thus, there are rarely simple, or even stationary, mappings between the action sequences executed and the homeostatic errors minimised.

In this paper, we briefly review theories and models providing an overarching framework as to how conjoint optimisation of physiological and behavioural homeostasis has been approached in neuroscience. We explore how this provides insight into the logic and provenance of primary rewards with respect to homeostasis. This prefaces an empirical section, where we review the neuroanatomical basis of glycaemic and energetic control in light of recent circuit-level evidence that illustrates the predictive nature of homeostatic control circuitry, and its putative role in modulating reward value computations. We will discuss how these homeostatic networks of the midbrain and brainstem innervate dopaminergic nuclei, and modulate reward (or precision) signals, in ways that are commensurate with their homeostatic and evolutionary imperatives.

## Theoretical accounts of homeostatic control

### Optimal control theory

Some of the earliest models of homeostatic control emerged from optimal control theory. Many of these were simple negative feedback systems where direct error correction was deployed to keep vital macro-state variables close to their set-points (Sterling 2012; Berridge & Robinson 2003). Common to most reactive schemes are the notions of a controller and a plant. The controller converts an input into a command which then inputs onto the plant, which outputs a motoric response, resulting in a new input. In the context of physiological regulation (Fig. 2a) the input to the controller is a homeostatic error, which is translated into a motor command. The command results in a behavioural exchange with the environment to reduce the homeostatic error. This new physiological state serves as the next input, generating the next homeostatic error. This feedback control gives rise to iterative error correction. In the case of glycaemic control, the homeostatic error would be inferred from the difference between current states (probed through central and peripheral glucose sensors) and a euglycaemic reference state (set point). The error is translated by the controller into commands for the visceromotor plant and the somatic motor system. The former creates autonomic gluco-regulatory responses – and the latter initiates a behavioural response such as foraging and consummatory action to minimize the homeostatic error. See also Powers (2016) for treatment of control theory from the perceptual side; in other words, the notion that phenomena such as homoeostasis and allostasis can be cast purely in terms of keeping sensations within bounds.

**Figure 2.**
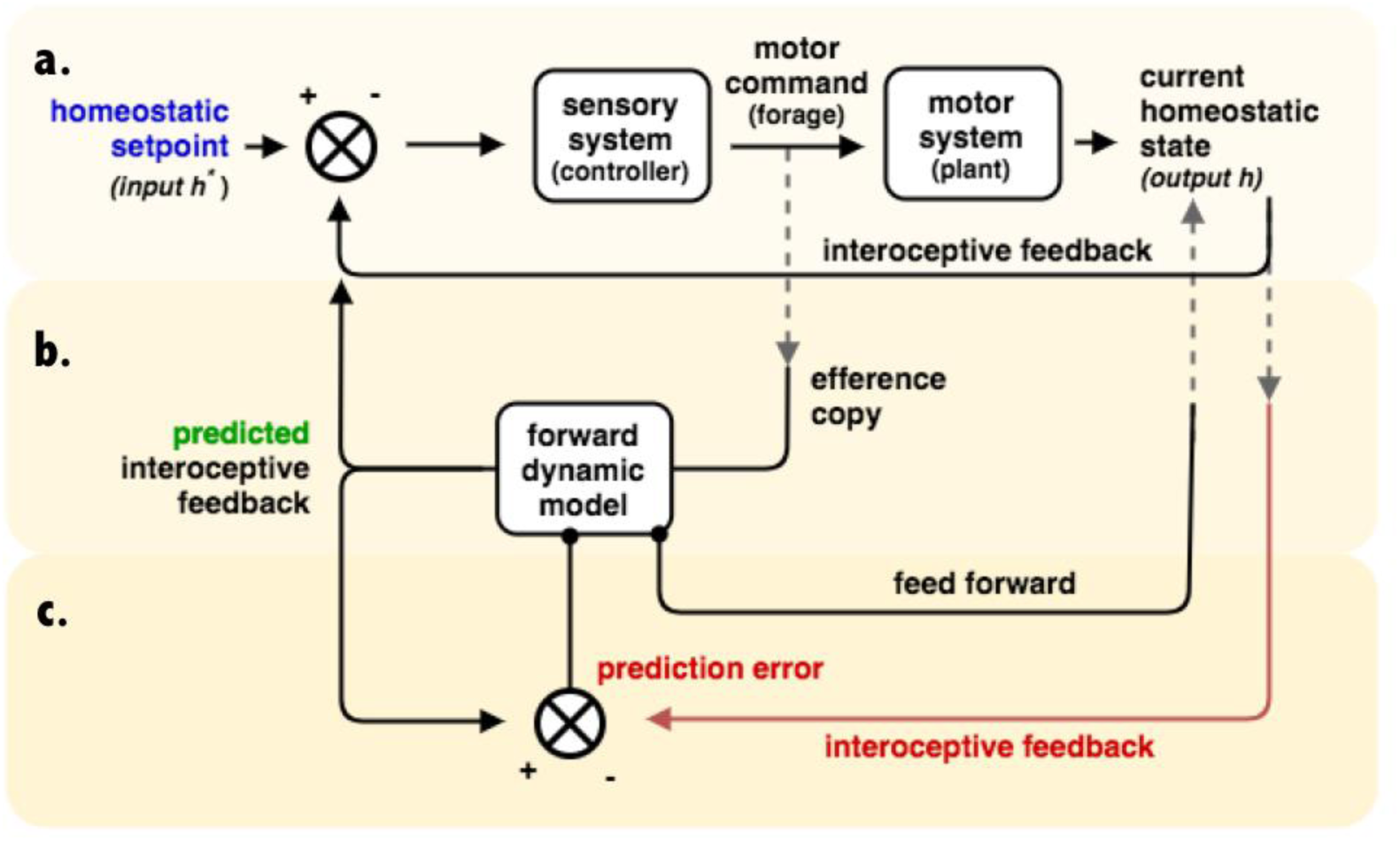
Comparator-based models. Under this class of model, the agent is described as a homeostatic error-correcting system. Broadly, the brain receives viscerosensory input from the body given its current physiological state, and it computes the homeostatic error between its current state and its set point (h*), and then iteratively deploys controlled action to correct this. **a**, Depicts the subsystem that entails direct feedback control, which combines a controller (here the sensory system), with a plant (here the motor system) that executes action to influence the current homeostatic state. The current homeostatic state is sensed by interoceptive feedback, which when compared to homeostatic set point, results in a homeostatic error that is forwarded to the controller, from which further motor commands are sent to the plant to iteratively minimise error. This is the homeostatic mechanism described by most physiology textbooks. **b**, An efference copy of the motor command is sent to a forward dynamic model that predicts the future interoceptive feedback, given the motor command. Residual errors between the predicted state and the set point are then iteratively minimised with further commands. **c**, Finally, a prediction error, computed as the error between the predicted and the current state is used to update the forward dynamic model. Insofar the system minimises this prediction error, the forward dynamic model makes accurate predictions of the homeostatic consequences of its actions.

One problem with direct feedback systems is that they are noisy and unstable (Carpenter 2004). Delays or noise in the control system can lead to error hunting, which results in oscillatory error corrections around the set-point. One solution is to introduce a predictive component (a forward model) into the control loop. In addition to an output command for the plant, direct feedback control, the controller generates an efference copy that is sent in parallel to the forward model (Fig. 2b). The forward model generates a prediction of the sensory state that is anticipated from the execution of the action, which feeds back and is compared to the set point. This culminates in a homeostatic prediction error that re-enters the controller and, if the forward model reliably predicts future homeostatic error, the controller acts accordingly to minimise this anticipated error. This logic can be expanded to include prediction error updates to the forward dynamic model (Fig. 2c).

Updating the forward model by discrepancies between predicted and actual interoceptive signals, is the fundamental feature of predictive processing and can (in principle) offer scope for explaining predictive homeostatic control outside the domain of purely reactive schemes. Such predictive control structures have been deployed in neuroscience to explain many phenomena, including motor control (Miall & Wolpert 1996) and awareness (Frith 2012). For comparison with the other theories we introduce below, we summarise the comparator based models of control theory in the upper part of Fig. 4.

Another feature that can be added to any of the models above is integral feedback control, where the time-integral of the error is controlled (Astrom 1995), rather than just the current or projected point estimate. Integral feedback control tracks a steady-state condition and only performs its regulatory action when this steady-state is violated. This form of control ensures that the system variable (e.g. glucose) returns back to the set point after a sustained step change irrespective of its magnitude. Integral feedback control account for the control of chemotaxis in bacteria (Yi et al. 2000; Barkai & Leibler 1997) and in systems neuroscience to explain flexibility of arousal and inhibitory control of the hypothalamus (Kosse & Burdakov 2014).

The models discussed above are deterministic in the sense that they operate under the assumption that the controller is already equipped with homeostatically rational commands to issue under the spectrum of hierarchically organised errors it can receive. In other words, such models do not by themselves offer any solution to the difficult problem of behavioural homeostasis in an uncertain and volatile environment. Answers to questions of the sort “*Which sequence of actions should I perform if I want to minimise this homeostatic error?*” are not addressed. If one is seeking to account for the conjoint optimisation of behavioural and physiological homeostasis, this is a serious limitation. Another limitation of this class of model is that it provides no principled means as to how to arbitrate between commands that entail different bundles of homeostatic error reductions; say between minimising one unit of thermal error (e.g. 1°C) and 2 units of osmolality error (e.g. 2 mOm/kg), versus 3 and 1 units, respectively. A seemingly sensible solution is to compute an aggregate homeostatic error as the Euclidean distance from set point, and choose the action that minimises that error. However, simply changing the units of measurement (e.g. from Celsius to Fahrenheit) inherently imposes an arbitrary prioritisation of one homeostatic dimension over another. The aforementioned feedback control models lack any discernible principles which would allow for such prioritisation to be achieved in any biologically meaningful way.

### Drive reduction theory

The problem of coordinating and prioritising multiple homeostatic feedback processes was a major inspiration to one of the earliest and most ambitious attempts at modelling behavioural homeostasis; namely, drive reduction theory (Hull 1943). Drive reduction theory was the first theory to algorithmically tether negative feedback to homeostasis via motivational drive. Instead of direct feedback via single homeostatic variables, motivational drive was proposed as a superordinate internal variable that is to be minimised over the long-run.

Under drive reduction theory, drive compels biological agents toward actions that remediate the basic physiological needs, in order to promote survival: “*…when any of the commodities or conditions necessary for individual or species survival are lacking, or when they deviate materially from the optimum, a state of primary need is said to exist*.” (Hull 1943). Drive can thus be conceptualized as a negatively valenced state that the agent works to attenuate. In so doing, the agent attenuates the associated homeostatic deficits that cause it. Stimulus-response associations are reinforced as a function of the resulting drive reduction – a postulate refined from Thorndike (1927). The reinforcement that accumulates over time determines the strength of habitually generating a response to a given stimulus (i.e., habit strength). The probability of executing a given action (i.e., the reaction potential) is determined by both habit strength and drive. More complex formulations take into account the inhibitory effect of fatigue, but the logic is the same. Drive-reducing actions are reinforced into habits, a behavioural means by which to minimise homeostatic error.

While drive reduction theory provides an integrated account of how deviations from the homeostatic optimum motivates behaviour, the theory falls short of explaining anticipatory behaviour that precedes any change in motivational drive. Animals develop drive states prior to any observable homeostatic deficits such as eating when sated, drinking before blood osmolality dips, and shivering before the onset of thermal challenges (Brown 1953; Sheffield & Roby 1950; Seward 1956; Bolles 1968). These early experiments show that the mechanistic account of drive reduction theory on learning is poorly predictive of behaviour, even in narrow experimental conditions.

### Homeostatic reinforcement learning

Reinforcement learning, a branch of machine learning inspired in part by behavioural psychology (and optimal control theory), offers some advance on the problem of homeostatic control. In any environment endowed with temporal regularities between sensory cues, actions, and outcomes, agents maximise expected future reward through algorithms that enable anticipatory action. The overarching aim of the agent under reinforcement learning is to maximize cumulative reward over some temporal horizon (Sutton & Barto 1998).

This can be achieved with several algorithms, and the dominating perspective on the computational role of dopamine in behavioural motivation stems from one such algorithm; namely the temporal difference (TD) algorithm. TD-learning relies on the difference between temporally sequential estimates (or predictions) of reward. If the prediction is wrong, the difference between previously predicted return (rational expectation of discounted rewards) and the new predicted return (predicated on the outcome observed) is computed as a prediction error, which is used to update the future prediction (much like in the above description of predictive processing). This is the foundation of the reward prediction error hypothesis (RPE), which states that phasic firing of dopaminergic neurons in the ventral tegmental area (VTA) and substantia nigra (SN) encode a reward prediction error signal (Montague et al. 1996). This theoretical prediction was later experimentally corroborated (Schultz et al. 1997), and since then much experimental work has underscored the importance of reward prediction errors in neurobiological accounts of learning and decision-making (Glimcher 2010; Niv et al. 2005). Several models have formulated homeostatic control using reinforcement learning algorithms (Dranias et al. 2008; Keramati & Gutkin 2014). With respect to homeostatic control, the perspective offered by Homeostatic Reinforcement Learning (HRL, Fig. 3) is interesting as it tessellates the core idea of drive reduction as sketched above, with reinforcement learning (Keramati & Gutkin 2014).

**Figure 3.**
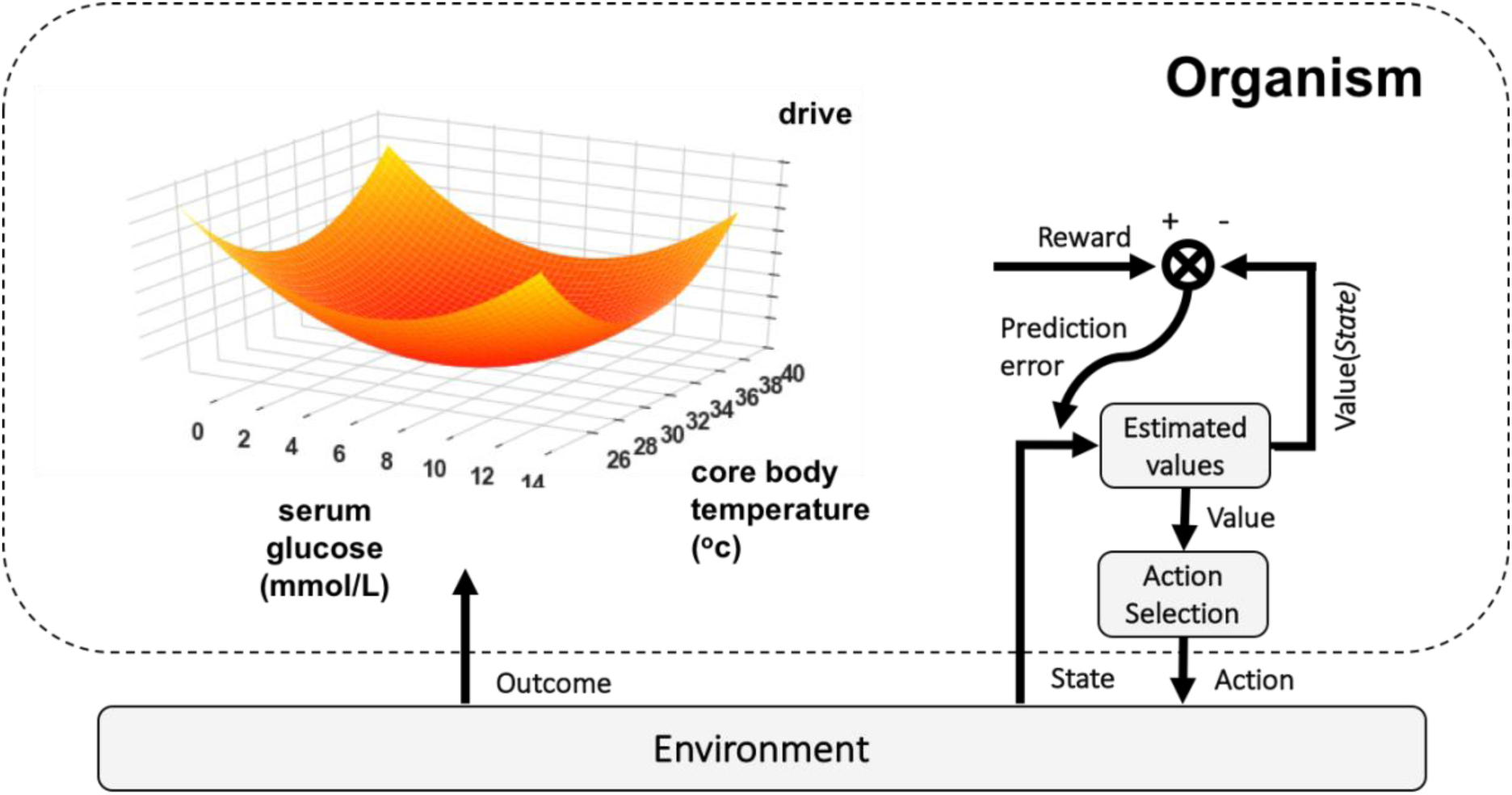
Homeostatic reinforcement learning. In the upper left, the surface represents a drive function, mapping from homeostatic state space (horizontal plane) to drive (vertical axis). The drive function depends on the homeostatic state space, and the system to be modelled. Here, we illustrate a drive function based on the surprise (negative log probability) derived from the survival probability function illustrated in Fig. 1b. If the drive function is appropriately configured, then actions – that influence homeostatic state such that homeostatic error is reduced – result in drive reduction. Under HRL, drive reduction is defined as rewarding, as in drive reduction theory. By comparing the estimated value to the actual reward experienced (with negative reward as drive inflation), a reward prediction error is generated and used to update future estimates of value. Actions are selected as a function of these estimated values, such that selecting the actions that maximise value, result in environmental exchanges that minimise drive, maximise reward, and thus minimise homeostatic error. Adapted from (Keramati & Gutkin 2014) with permission.

The HRL framework defines a homeostatic state space, from which a drive function is derived, mapping non-linearly from homeostatic state to drive (Fig. 3, & 4 middle). The central logic is that with drive reductions defined as reward, agents that learn to maximise reward, will minimise drive, which minimises homeostatic error, meaning that reward maximisation and homeostatic regulation (behavioural homeostasis) are “*two sides of the same coin*” (Keramati & Gutkin 2014). HRL accounts for anticipatory features of behavioural homeostatic control, showing that simulated agents could learn to incur short-term homeostatic errors (e.g. deviations from a set point), in order to mitigate long-run (path integrals of) homeostatic errors. While the HRL framework accommodates anticipatory behaviour of homeostatic control, it is worth pointing out some of the residual problems. Strictly speaking, HRL theory specifies no criterion to define the biological maximandum (i.e., the optimal set point), but relies on experimenter-set value functions which have no normative grounding. Keramati and Gutkin (2014) choose their drive function as a sensible and parsimonious guess based on the behavioural and economic phenomena this would entail. Interestingly they showed that several phenomena from economics and behavioural ecology could be accounted for with a simple convex drive function. To facilitate comparison between theories, HRL is juxtaposed with other models in Figure 4.

**Figure 4.**
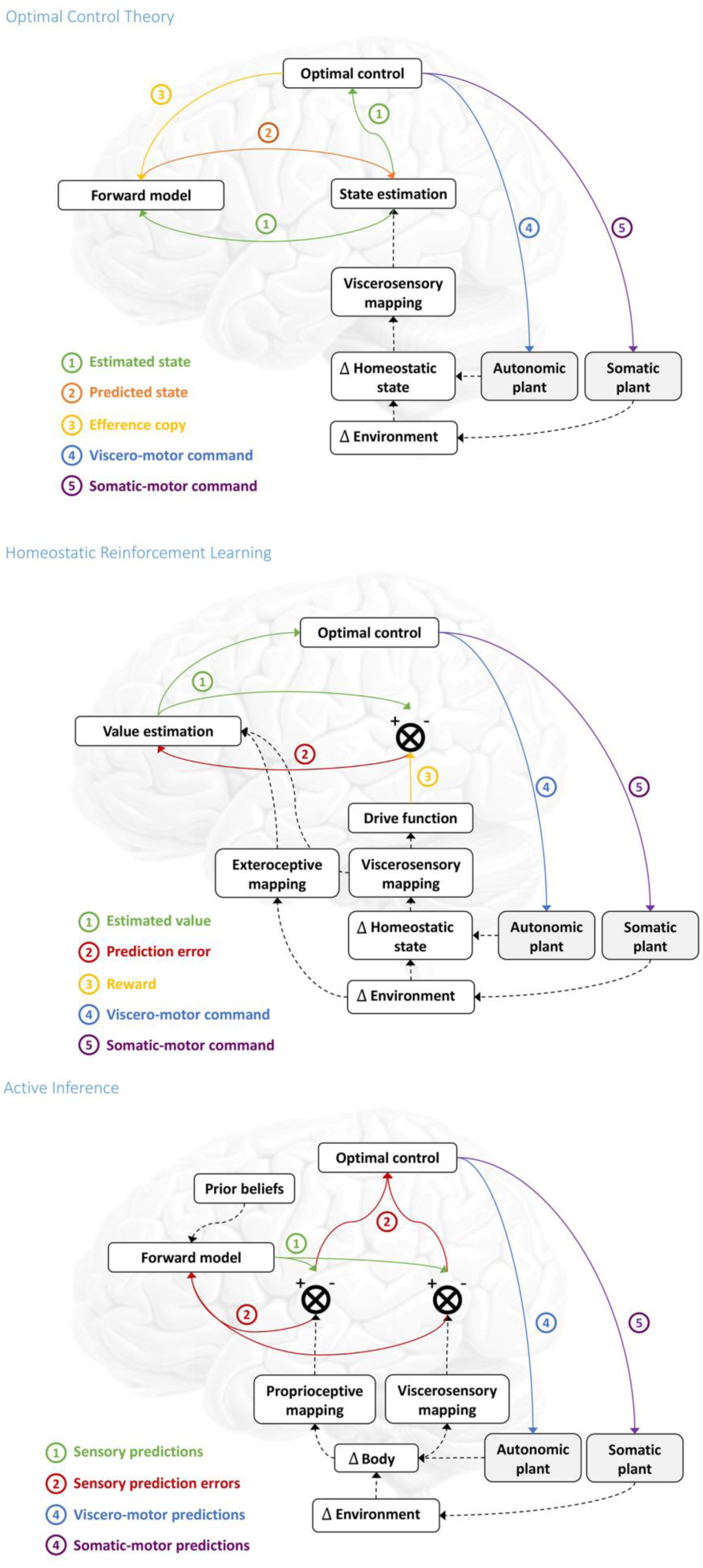
**Comparing models**. Common to all model architectures is the fact that the agent is in a given homeostatic state and a given environmental state. The agent can act in two ways. First it can engage in physiological homeostasis by controlling its autonomic plant, which directly modulates the homeostatic state of the body (∆homeostatic state). Second it can engage in behavioural homeostasis by moving, via the somatic-motor system, to change its sampling of the environment (∆environment), which will indirectly change its homeostatic state. These changes in homeostatic state are hidden, but they can be transformed into neural input by the viscerosensory system (viscerosensory mapping). How the different theories prescribe the control of these two plants based on sensory inputs is highlighted in the following comparison. **Optimal control theory**. This is a schematic summary of the components commonly found in conventional treatments of optimal motor control, here applied to homeostatic control. The hidden homeostatic states produce interoceptive sensations through the viscerosensory mapping. This viscerosensory input is used for hidden-state estimation (e.g. by Bayesian filtering) based on the forward model and a (weighted) prediction error. The prediction error is the difference between sensory input and predictions of that input given the predicted state (orange). The state estimates are used for optimal control, which returns motor commands (purple and blue) that minimize future cost or loss, specified by a cost function (not shown, alternatively known as an inverse model). These optimal control signals are then sent to the two motor plants and (through an efference copy, yellow) to the forward model. The forward model then computes the predicted change in hidden homeostatic states. In this scheme, the forward model can be regarded as a mapping from motor control to changes in hidden homeostatic states. Effectively, its role is to finesse the problem of inferring homeostatic states and thereby optimize homeostatic control signals. This is necessary because delays and noise on sensory signals could easily confound the implicit closed-loop control used by this scheme. **Homeostatic reinforcement learning**. Here we interpret the schematic in Fig. 3 using the same logic and terms wherever possible. Again, we start with the hidden homeostatic states that are sensed via a viscerosensory mapping. This viscerosensory input is submitted to a drive function, mapping from sensory state to the negative valenced motivational drive. The drive reductions are encoded as experienced rewards (maroon), which are subject to a temporal difference learning, where the value of sensory states are estimated as the rational expectations of future discounted rewards following from that state. The difference between the value estimated at a given trial (green), and the updates to that value based on the new sensory inputs (exteroceptive and viscerosensory), yields a reward prediction error (red), that is used to update the value estimate. The value estimates (green) for different possible actions, are then subject to action selection, from which the highest value action can be probabilistically selected. **Active inference**. We start again by considering environmental dynamics caused by somatic action. Along with autonomic action, this can result in changes to the body, causing both proprioceptive and viscerosensory input (we omit exteroceptive sensations for clarity), yielding proprioceptive and viscerosensory prediction errors. These prediction errors are simply the difference between the sensory input observed and the sensory inputs predicted under the predicted (hidden) states. In this form, top-down predictions from the forward model are compared with sensory inputs to produce bottom-up prediction errors (red connections) that enter the forward model. Crucially, the mapping from hidden states to sensations is now part of the forward (and thus, generative) model. Here, cost functions have been replaced by prior beliefs about (desired) homeostatic states. Allostatic regulation here can be achieved through prior beliefs over homeostatic trajectories. These prior beliefs enter the forward model to guide predictions of sensory inputs. These prior beliefs set the targets and priorities of homeostatic control, and thus are strongly selected as a function of their contribution to survival (and thus fitness). Proprioceptive predictions are fulfilled by the somatic motor system by classical motor reflex arcs (the somatic plant), while predictions of viscerosensory input are fulfilled by the autonomic plant. Optimal control now reduces to simply suppressing proprioceptive and viscerosensory prediction errors.

### Active inference

Recent models invoke the notion of variational inference under a hierarchical Bayesian model to solve homeostatic control problems (Stephan et al. 2016; Pezzulo et al. 2015). Fundamental to those formulations is the notion that the agent deploys interoception, somatic and viscero-motor actions in order to control internal states. This is framed under active inference (Fig. 4, lower), which is a corollary of the free energy principle (Friston et al. 2006; Friston 2012). Heuristically, this principle suggests that all living agents resist disorder (i.e. death) by restricting themselves to a limited number of states consistent with their physiological integrity, an idea that is consistent with homeostatic regulation as framed above, and with drive reduction theory.

Under active inference, agents stay alive by predicting the states that keep them alive, and act in order to fulfil those predictions. These predictions are generated in the higher levels of the neural and autonomic hierarchies and passed down to lower levels. The lower levels signal prediction errors back up the hierarchy. Prediction errors here are not about reward per se, but rather discrepancies between expected and realised sensory input. Sensory predictions are cascaded downwards in the hierarchy, and if it does not match input, prediction errors are propagated upwards in order to update the model (interoception) or act on the environment in order to change the sensory input via (motor and autonomic) reflexes (Fig. 4, lower). Importantly, agents are endowed with prior beliefs that are congruent with high-survival states, such as being sated, hydrated and warm. As such, the notion of reward – common to models of reinforcement learning and optimal control – is absorbed into expectations about occupying states that increase biological fitness. Any action that underwrites the probability of fulfilling those expectations can be said to have value.

It is useful here to compare and contrast control theoretic formulations with active inference in the proprioceptive domain, because the same principles may apply in the interoceptive domain too. In the control of striated muscle, active inference formulations of motor control replace motor commands with predictions of proprioceptive sensations. These predictions afford the equilibrium or set points that enslave classical motor reflexes or goal-directed actions. This control architecture calls upon earlier notions such as the equilibrium point hypothesis (Feldman 1986), in which desired movements are specified in terms of equilibrium or fixed points. Clearly, as above, the question now arises: Where do the predictions or equilibria come from? In active inference, these are generated by a deep (generative) model that provides contextualised predictions that are fit for purpose, in the current context (Friston et al. 2017). In other words, hierarchically high level motor goals specify predictions of subgoals and so on – all the way down to the predicted primary sensory afferent input in the spinal cord or brain stem. The crucial aspect of this architecture is that the forward model is not used to nuance feedback control (as in comparator models of optimal control theory, e.g. Fig. 2 & Fig. 4 upper) – it plays a foundational role in prescribing behaviour as a generative model (Fig. 4, lower). Furthermore, this architecture is effectively open loop because its set points are predefined by descending predictions. However, these predictions are generated from a hierarchical synthesis that contextualises them; rendering the overall system a closed loop architecture. The argument in this paper is that exactly the same mechanisms apply in the context of homoeostasis through allostatic responses that rest upon purposeful behaviour in response to the interoceptive and exteroceptive cues.

Ultimately, action and interoception serves to fulfil predictions of homeostatic equilibria on all levels of the hierarchy, from autonomous physiological processes to behavioural homeostasis: Autonomous processes, such as the release of insulin from the pancreas when glucose levels drop, most likely constitute the lower layers in the hierarchy of the homeostatic reflex arc and are most likely implemented by effector regions in the spinal cord and brainstem (Seth 2013; Stephan et al. 2016). Premeditated planning and decision-making that engenders allostatic change is governed by relatively higher layers in the control structure, e.g. in the prefrontal, insular, or anterior cingulate cortex (Stephan et al. 2016). Thus, the hierarchical structure of models suggested under active inference, has the potential to account for homeostatic regulation to unfold on all spatiotemporal scales relevant for physiological and behavioural homeostasis. It is the hierarchical architecture implicit in active inference that accommodates the spectrum of spatial temporal scales; providing a hierarchal distinction between high level predictions (allostasis) and low level predictions (classical homoeostasis). In this setting, low-level interoceptive prediction errors that cannot be resolved immediately are passed to higher levels to induce deliberative behaviour that, in the long-term, returns physiology to its fixed (set) points.

A central concept for active inference accounts of homeostatic control is the notion of information theoretical (Shannon) surprise. Technically, surprise is the negative log probability of a state – which coheres with the intuition that an internal state that is highly probable – carries less surprise than one which is improbable. Importantly the level of surprise scores how valuable states are, since the most probable states (the low surprise of occupying internal states close to set point) are most probable because they afford the highest probability of survival, whereas the least probable states (the high surprise of occupying extreme internal states) are the least probable because the afford the lowest probabilities of survival. The high surprise states are thus the states in the tails of the survival probability surface in Fig. 1b. This closely relates to another concept from information theory; namely, entropy, which is simply average surprise. The overarching aim of the adaptive agent is to keep sampling sensory data that is as unsurprising as possible, because the agent expects to constantly find itself in homeostatic equilibria, minimising its entropy. This prior belief (of being close to a set point) is engendered by a generative (forward) model, yet another key concept in active inference, to which we now turn.

A generative model establishes a probabilistic map between hidden causes (internal or external states) to observed consequences (proprioceptive, exteroceptive or interoceptive sensory input) by combining a prior (here, encoding the prior probability of internal states) with a likelihood function (a probabilistic map from hidden internal states to observed sensory inputs, see Fig. 5). Principally, there are two means by which prediction error and thus surprise can be minimised. The agent can update its predictions to conform to the sensory input (interoception), or act on the world to change the sensory input generated by external states, to better match its predictions (action). The interested reader should see Bogacz (2015) for an tutorial based introduction to the technical aspects variational inference in context of perception.

**Figure 5.**
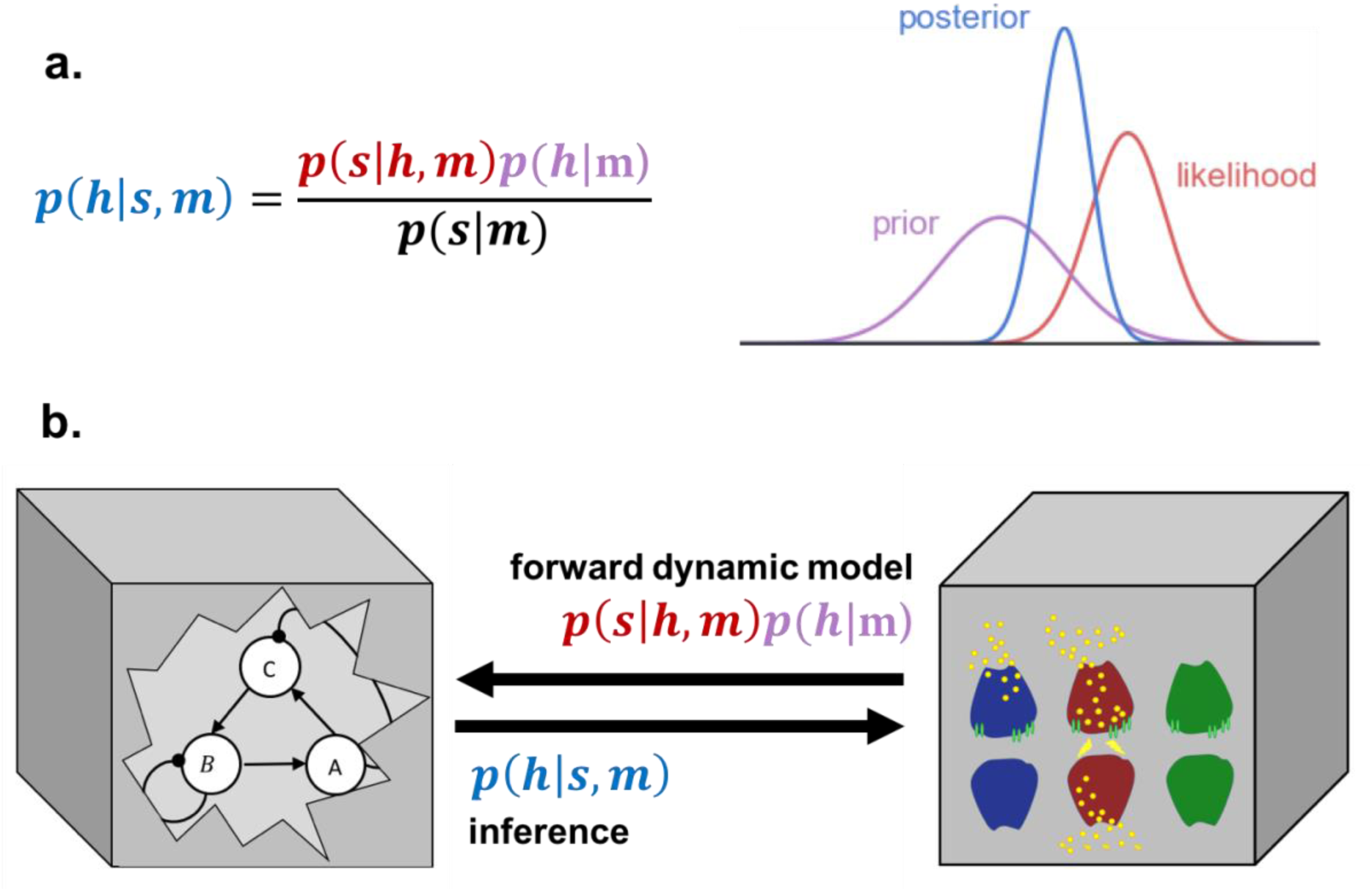
Interoception, Bayes rule and generative models. **a**, Bayes rule provides the probabilistic foundation for the generative model *m*, which the agent embodies, and under which the agent expects homeostasis. This consists of a likelihood function *p*(*s*|*h*, *m*) (a probabilistic map from hidden external states, *h*, to sensory inputs *s*) in conjunction with a prior *(h|m)* (a probability distribution over external states, including bodily states). In this setting, the prior can be interpreted as a probabilistic set point. Calculating the posterior *p*(*h*|*s*, *m*) is model inversion and yields the probability of a hidden homeostatic state, given the sensory input. Thus, the posterior is a mix between likelihood (how likely is this) and prior (how often does it occur) weighted by their relative precision (see Bogacz 2015). The denominator *p*(*s*|*m*) is a normalisation term that ensures the posterior integrates to one. Importantly, this term is also the foundation of Bayesian model selection (see Ghahramani 2012 for an introduction). **b**, Illustrates how homeostatic dynamics (left box) that are hidden from the agent give rise to sensory input *s*, which the phenotype must infer on, given its generative model *m*. The causal structure of the external world (including the body) is encoded in synaptic activity (right box) encoded in a forward dynamic model, which allows inference about the causes (hidden states) of the sensory input.

When considering homeostatic control as active inference, it is important to appreciate the nature of prior beliefs. In a hierarchical setting, these are referred to as *empirical* priors. This is because they can be informed by empirical data or sensations. This leads to a picture of the interoceptive hierarchy as encoding a cascade of prior expectations and subsequent predictions for the level below. In most formulations, deeper (i.e. higher) expectations usually entail longer time courses or horizons, while priors at lower levels are more concerned with proximal outcomes. On this view, surprising violations (i.e., departures from homeostatic set points) induce ascending prediction errors throughout the hierarchy until some (allostatic) expectations change the organism’s circumstances. Under this framework, it is likely that some empirical priors are held with greater precision (e.g. body temperature), and thus prevail with only minor modulation over many different settings, while others will be lower in precision, and thus have greater latitude to be informed by context (e.g. hunger). We will see later, that the precisions – afforded different prediction errors at different levels of the hierarchy – are a key determinant of behaviour and the balance between allostasis and classical homoeostasis.

In short, prior interoceptive beliefs should reflect (relatively) invariant survival probabilities, and should only be (allostatically) modulated to reflect a shift in the peak survival probabilities. A good example of this would be having a relatively invariant prior belief about what core thermal states the agent should occupy, but then modulating this under conditions of viral infection, where the survival probability function shifts such that higher thermal states have the highest survival probabilities; hence the phenomena of fever, and its related thermoregulatory behaviours.

Prior beliefs about homeostatic set points are likely to be hardwired in effector regions, such as the hypothalamus and brainstem nuclei. Such empirical priors are likely to be genetically specified and shaped via evolution as a function of their ability to minimise surprise, given the agents respective eco-niche (Friston & Ao 2012). On the other hand, priors that pertain to learning and adaption must be able to change during interaction with a dynamic, hierarchical and often volatile environment. For a more expansive account of learning and homeostasis under active inference see Pezzulo et al. (2015).

### Summary

In the above we framed the problem of homeostasis, not as a problem of stability per se, but rather as predictive control over the physiological and behavioural processes that keep vital homeostatic variables within the narrow (but dynamic) range that ensure survival. We rehearsed some early attempts at modelling such control, using various schemes of feedback control. While these may suffice for physiological homeostasis through autonomous control (e.g. the baro-reflex or skeletal muscle control) they are often unstable, and importantly do not afford any insight into the mechanisms of behavioural homeostasis that unfold over longer timescales. Reinforcement learning solves this shortcoming by proposing several algorithms that frame adaptive behaviour as reward maximisation, which can be harnessed to defend a homeostatic set point (Dranias et al. 2008; Keramati & Gutkin 2014). One exigent problem (see Friston & Ao 2012 for several others) with reinforcement learning in general is that the definition of reward is behaviour-centric: Agents strive to maximise reward, but reward is defined from observed behaviour. Or as Berridge (2004) puts it “*A circular explanation is one that attempts to explain an observation in terms of itself. It just reasserts what has been observed and does not really add any new explanation*.” Avoiding this circularity through homeostatic considerations was a central motivation for the development of Homeostatic Reinforcement Learning. Likewise active inference accounts of adaptive behaviour avoid this circularity by providing a normative account of why agents must necessarily infer and minimise surprise about their own internal hidden states in order to maintain physiological integrity (Friston 2012; Friston et al. 2006). This hierarchical Bayesian perspective absorbs the entire suite of concepts discussed above (see Stephan et al. 2016 for details). Concisely, set points and error functions that are integral to any form of feedback control are replaced by prior beliefs (or predictions) about sensory input, where subsequent deviation from those beliefs is encoded as the errors of prediction (Fig. 4 lower).

Furthermore, the conceptual objects of reward and value that motivate behaviour (as defined in reinforcement learning), are absorbed into prior beliefs about the consequences of action (e.g. what actions minimise prediction errors), where desirable outcomes are simply those that engender the least surprising outcomes. So far, we have discussed active inference in general terms; in a way that places the predictions of hierarchal or deep generative models centre stage. To properly understand the implicit computational architecture that underwrites allostatic responses, it is worthwhile unpacking the imperatives for active inference in terms of *resolving uncertainty*. Formally, uncertainty is expected surprise. Therefore, to select policies that minimise expected surprise in the future, one has to evaluate the associated uncertainty in terms of *expected free energy*. Expected free energy usefully decomposes into epistemic and pragmatic terms – usually associated with intrinsically motivated, information-seeking, epistemic behaviour on the one hand and extrinsically motivated, reward-seeking, pragmatic behaviour on the other. The epistemic part is important for allostatic responses (and is generally ignored in reinforcement learning formulations). A simple example here is the epistemic value or affordance of checking whether the fridge is contains the necessary ingredients, before starting to prepare a meal, or the foraging mammal scanning its environment to infer the location and habits of its prey. Typically, uncertainty reducing (expected free energy minimising) policies are selected that first resolve uncertainty after which, prior preferences come to dominate. This leads to a structured transition from explorative to exploitative behaviour. They can also be selected under satiety states, where homeostatic errors are attenuated, and the value of exploitative action is diminished.

One subtle aspect of this construction is that we now need to posit generative models that entertain the future consequences of action. Although obvious, this means that there must be neuronal representations of (worldly and bodily) states in the future, under each competing policies. These counterfactual futures may have limited time horizons, but must exist under the theory. The resulting deep generative models are sometimes referred to as having *counterfactual depth* that necessarily entails a future. The notion of counterfactual encoding (i.e., neuronal representations of future states) is therefore something that should figure, when trying to understand interoception and its role in homoeostasis (Seth 2014).

Crucial for our argument is that policy selection depends upon the degree to which a given policy will resolve uncertainty and the confidence or precision placed in the ensuing beliefs about policies. In other words, to select the best policy, one has to evaluate the precision or confidence in beliefs about alternative ways forward. A body of evidence now points to dopamine as signalling fluctuations in the precision or confidence associated with policy selection (Fiorillo, Tobler et al. 2003, Niv, Duff et al. 2005, Humphries, Khamassi et al. 2012, Friston, Schwartenbeck et al. 2014, Schwartenbeck, FitzGerald et al. 2015). This will become relevant later when we interrogate the empirical evidence that speaks to different theoretical formulations of homoeostatic control.

## Neural bases of energetic control

In the following empirical section, we survey recent evidence that suggests that particular circuits of the hypothalamus and brainstem play a role in predictive homeostatic control. We will focus exclusively on energetic control, as the experimental evidence for this homeostatic dimension is extensive and (at least relative to other homeostatic dimensions) easy to manipulate and measure. This subfield also contextualises the common use of hunger as the predominant motivational strategy for animal experiments.

### Hypothalamus as a homeostatic controller

Situated inferior to the thalamus and superior to the pituitary gland, the hypothalamus is an archipelago of distinct nuclei, charged with coordinating a microcosm of homeostatic functions (Fig. 6a). The existence of opponent energy-regulating processes was an early and exciting discovery; two hypothalamic regions with opposing effects on food intake were found, a lateral area resulting in hyperphagia when stimulated (‘feeding centre’), and a ventromedial area resulting in hyperphagia when ablated (‘satiety centre’, Aand & Brobeck 1951; Brobeck 1946). Since then, modern cell-type specific techniques for circuit manipulation and projection-specific has afforded an unprecedented window into the deep and neuroanatomically complex networks involved in energy homeostasis. One of the major components of these networks is the arcuate nucleus (ARC), lying in the mediobasal hypothalamus, on either side of the third ventricle, just above the median eminence. There also, at a finer sub-nuclei scale, opponency remains an important principle. Two cell types are found to be crucial for the control of feeding (Atasoy et al. 2012), identified by expression of the neuropeptides Agouti-related Protein (AgRP) and Proopiomelanocortin (POMC), which have seemingly opposing properties.

**Figure 6.**
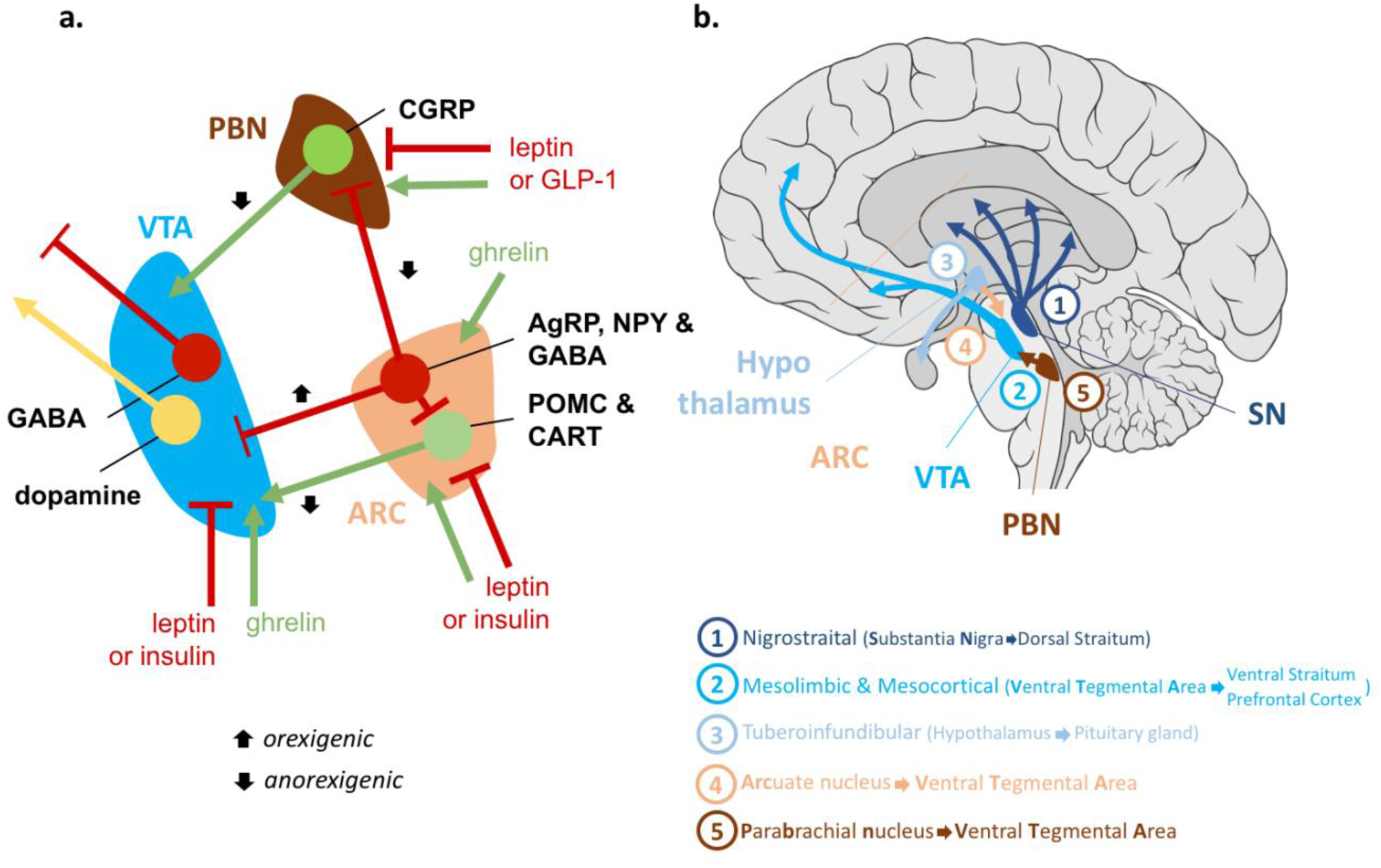
Homeostatic-reinforcement interface. **a**, Red T-lines and lettering illustrate inhibitory inputs; Green arrows and text indicate excitatory inputs. Dopamine is coloured yellow as this can have both excitatory and inhibitory effects depending on receptor subtypes. Projections: Cocaine and amphetamine regulated transcript (CART) and pro-opiomelanocortin (POMC) in the ARC of the hypothalamus process POMC to alpha-MSH that activate melanocortin-4 receptors (MC4R) on post-synaptic cells in the arcuate of the lateral hypothalamus (ARC), which projects to the VTA (details not shown here, see Ferrario et al. 2016). This melanocortin pathway is suppressed by neighbouring cells in the ARC that produce Agouti-related protein (AgRP), neuropeptide Y (NPY) and GABA that all inhibit POMC neurons (Mandelblat-Cerf et al. 2015) and project to many of the same sites, including the VTA. Further, these project to CGRP neurons in the parabrachial nuclei, which in turn projects to VTA. Hormone input: AgRP neurons are inhibited (red T-bar) by leptin and insulin, whereas POMC are activated by those same hormones (green arrow). Hormone ghrelin, that signal short term energy deficits, activates AgRP and dopamine (green arrows) in the VTA (Palmiter 2007). Conversely, leptin and insulin attenuates dopaminergic firing (red T-bar). **b**,. There are four important dopaminergic pathways that project from the midbrain widely through the brain. Importantly, the VTA projects through the mesolimbic & mesocortical reward circuit to the caudate & nucleus accumbens (NAc) in the striatum and also the amygdala, hippocampus and prefrontal cortex. Further, the VTA also hosts GABAergic projection neurons that modulate many of the same target regions as dopamine.

AgRP neurons are activated by energy deficits (Mandelblat-Cerf et al. 2015), report on the nutritional state of the body, and are both necessary (Luquet et al. 2005) and sufficient (Aponte et al. 2011) to evoke voracious feeding and food-seeking behaviours: the number of stimulated AgRP neurons is linearly predictive of food intake. These effects appear to be mediated by GABA and the neuropeptides NPY and AgRP, that stimulate food intake when delivered directly to the arcuate nucleus. In the absence of food, stimulation of AgRP neurons promote a range of learned behaviours that relate to hunger and food-seeking (Dietrich et al. 2015). POMC neurons by contrast are activated by energy surfeit and their activity inhibits food intake and promotes weight loss (Atasoy et al. 2012). AgRP and POMC neurons are both regulated by the circulating endocrine signals of nutritional state, modulating their activity in mutually opposing directions consistent with their function. These two cell types interact in part through a common set of downstream melanocortin expressing neurons that are activated by POMC and inhibited by AgRP. These two subpopulations are interspersed within the ARC making it an obvious candidate site for the encoding prediction errors for energetic wealth (energy balance, or other synonyms).

### Predictive responding

A recent stream of research has employed cell-specific techniques to image and causally manipulate the activity of AgRP neurons under different homeostatic challenges that each manipulate homeostatic error, and thus causally control the motivational state of the animal. Natural deprivation, ghrelin injection, pharmacological or optogenetic activation of AgRP neurons evoke voracious feeding and inhibit POMC neurons, as might be expected with a large deviation from a set point (Betley et al. 2015; Chen et al. 2016; Krashes et al. 2014; Mandelblat-Cerf et al. 2015). However, a homeostatic-comparator based view of the hypothalamus has been challenged by several recent papers that show AgRP and POMC neurons encode predictive signals, varying as a function of future expectations, rather than currently realised energy states per se.

### The sensory paradox

These aforementioned papers show for example that fasting-activated AgRP cells are inhibited by the visual presentation of food, prior to eating. This phenomena appears to be paradoxical (the sensory paradox, hence) for the homeostatic view of AgRP encoding drive (Wise 2013). If AgRP neurons encode feeding or drive (hunger) how can they switch off prior to feeding, given that drive is not immediately mitigated upon seeing the food? Emphasis on the surprising nature of this result, now replicated several times, hinged on the fact that inhibition occurs even before the food is tasted. Yet, we would argue that the predictive nature of the signal, does not rest on it occurring before the taste of the food, since even if it were time-locked to the taste at consumption, it would still be predictive insofar as no change in nutritive wealth is yet manifest. Indeed, any candidate drive or hunger signal that changes reliably to extero- or interoceptive cues is still a predictive signal with respect to the slow dynamics of the gastro-intestinal cascade (the cascade of physiological events that happen after ingestion). Arguably the energy content of a food is not fully appropriated until the post-absorptive phase. In this light, the sensory paradox is just as much a paradox for interoceptive responses time-locked to consumption (like taste or olfaction), as they are to the exteroceptive signals underpinning cue-learning (e.g. sight). Since these responses are not taken to be paradoxical, it could be said that the sensory paradox somewhat dissolves.

In all reported cases to date, most of the above-baseline activity of AgRP neurons was inhibited prior to feeding initiation (Chen et al. 2016; Betley et al. 2015; Mandelblat-Cerf et al. 2015; Chen & Knight 2015). The degree of inhibition has been shown to depend on food quality, caloric content (Chen et al. 2016), and even show a rebound back to original levels upon the experimenter rescinding food. These up and down modulations of AgRP that vary as a function of the agents beliefs, are mirrored by the hormonal signals of energetic status, that can also be considered endocrine predictors of future energetic wealth: Leptin, putatively signalling positive energetic wealth (Domingos et al. 2011), suppresses AgRP (Takahashi & R. D. Cone 2005; Fulton 2000; Betley et al. 2015); whereas ghrelin, putatively signalling its converse, excites. The inhibition appears sensitive to the appetitive affordance of food (Gibson 2001), such that food presentation in a closed container that allowed sight and smell of food but not consumption, had diminished inhibitory effects (Chen et al. 2016).

Indeed, compatible with the fact that feeding can always be disturbed at any point, residual AgRP firing persists throughout the consummatory period (Mandelblat-Cerf et al. 2015). Through the lens of drive reduction theory, AgRP inhibition thus appears to track the expected drive that fluctuates with incoming sensory evidence (and thus how this updates the brain’s generative model). These findings are all compatible on the interpretation that AgRP firing itself encodes counterfactual prediction errors over a spectrum of near-term temporal horizons. On this hypothesis, AgRP should be inhibited by any exteroceptive or interoceptive cue that predicts reductions in energetic drive, and excited by any such sensory cues that predict inflations of energetic drive. This expectation would plausibly be predicated on an accumulation of evidence integrating sensory modalities. One obvious prediction would be that the AgRP baseline firing effect should, with sufficient training, be quantitatively sensitive to the predictive probability of sensory cues in both directions, signalling expected decrements and increments in expected energetic prediction errors. It should be noted that these expected future energetic prediction errors are prediction errors over viscerosensory states associated with energetic wealth, that likely follow from gastrointestinal and adipose systems. The way that energetic wealth predictions are derived from these redundant signals will be an important next step toward understanding the AgRP encoding function.

It is interesting to note that the AgRP responses are heterogeneous in their temporal kinetics (Betley et al. 2015), in responding to food-predictive cues, with some showing a slow attenuation over time, and others faster. This suggests that the AgRP population as a whole encodes a distribution of energetic errors over a spectrum of temporal horizons.

### Valence signalling

Another interesting parallel, between this new wave of AgRP data and drive reduction theory as outlined above, is that both drive and AgRP carry negative valence, as well as the fact that reducing-drive and reducing-AgRP activity are imbued with positive valence. This is a subtle issue, and can cause some seemingly conflicting conclusions, with some groups reporting that AgRP carries negative valence (Betley et al. 2015), and others reporting its positive valence (Chen et al. 2016). The discrepancy can arguably be resolved in light of DR. Under DR (and therefore its cognate, HRL), drive is a negative valence signal, that agents work to minimise. Actions that reduce drive are rewarding which reveals why the attribution of valence to neural signals could easily be conflated. The key prediction is that if AgRP signals future drive (an error on the predicted energetic wealth), then AgRP stimulation, in the absence of any means of reducing drive (i.e. food), should be aversive since drive-inflations are costs (negative reward). Indeed, AgRP stimulation can condition place (and flavour) aversion (Betley et al. 2015). However in the presence of food, the drive reduction that follows AgRP stimulation should be larger and thus more rewarding, thus the reinforcing effect of AgRP stimulation should only occur in the presence of food, which is indeed what is observed (Chen et al. 2016). This is indeed a key distinction between the two opposing papers. One apparent problem with this model, is that mice fail to perform operant responses in order to shut off AgRP neuron activity (Betley et al. 2015; Chen et al. 2016); however, it is important to consider issues of credit assignment. Under natural conditions, a drive reduction such as that associated with AgRP silencing, in the absence of sensory food cues, can only be due to post-ingestive effects. This means that the food consumed minutes or hours previously will be assigned the credit for the drive-reduction caused by AgRP inhibition now, which predicts that recently performed operant actions should not necessarily be reinforced at all. Under the mouse’s generative model of the world, (again in the absence of sensory food cues) the drive reduction should most likely be caused by actions/sensory/gustatory events long before the operant action was performed. How easily mice could learn this long-range temporal contingency with overtraining though is an open question.

## Interface between reward prediction errors and glycaemic control

### Introducing dopamine

The catecholamine dopamine is synonymous with reinforcement, reward and motivation. Whilst the literature on dopamine is vast, we will restrict discussion to its putative role in glycaemic or energetic control as discussed above. It is well known that phasic signals in ventral tegmental area (VTA-DA), and thus dopamine release in the mesolimbic system, systematically scale with the nutritive value of oro-sensory events in monkeys, where reward magnitude is determined by the volume of nutrients consumed (Tobler et al. 2005; Stauffer et al. 2014; Ballard & Knutson 2009). In humans, there is evidence that post-prandial dopamine release is modulated by deprivation with dopamine binding decreasing more in response to consumption after fasting compared to non-fasting (Small et al. 2003). This echoes extant evidence from rats and mice that show increased dopaminergic release (as measured by dopamine metabolite 3,4-dihydroxyphenylacetic acid) at feeding after a period of starvation in the nucleus accumbens (NAc, McCullough & Salamone 1992; Radhakishun et al. 1988), medial prefrontal cortex (but not NAc, Carlson et al. 1987) and interestingly, the posterior hypothalamus (Heffner et al. 1980). Despite these findings and others, one of the curious features of the literature on phasic DA and reward is that animals are motivated by a homeostatic deficit such as thirst or hunger, and yet homeostatic states are rarely foregrounded in analyses of relevant modulators of reward signalling. One recent interesting exception to this is offered by Cone and colleagues, who present evidence for how sodium depletion can modulate RPE in the NAc of rats (J. J. Cone et al. 2016). By pairing sodium sated and depleted rats with conditioned and unconditioned stimuli, they found that phasic dopaminergic RPE signals can manifest independently of learning and are “*expressed as a function of their current* [homeostatic] *value to the organism*” (J. J. Cone et al. 2016, square brackets added).

Thus, on many grounds, homeostatic states should be potent modulators of these DA signals. As the animal plays its task for consumption of water or sugar-containing juice, its homeostatic deficits diminish, or are predicted to diminish, meaning that the value of those commodities should steadily decrease. Indeed, given the quantitative evidence for a relation between RPE and marginal utility (Stauffer et al. 2014), the fact that this is rarely tested or acknowledged (or for that matter controlled for) is surprising, given that the manipulation that makes the outcomes rewarding is continually being attenuated, until the animal rejects further play, presumably because the marginal utility of consumption has depleted to a point of indifference. For this reason, we recommend greater scrutiny of homeostatic states, and their dynamics under neurobiological studies of reward. In the case of energetic variables, intra-arterial telemetric glucose monitors are now available, and could afford important insights in this regard.

At this point, we introduce a fundamentally different perspective on the role of dopamine. In schemes that commit themselves to some form of reinforcement learning, dopamine is usually cast as a reward prediction error (Fig. 4, middle); namely, the difference between expected and encountered reward. This is in contrast to active inference formulations, which accommodates the fact that dopamine is a neuromodulator. In other words, dopamine cannot drive synaptic responses – it can only modulate them. This modulatory role is exactly that required of precision. On this view, phasic dopamine responses signal an increase in the precision or confidence placed in beliefs about ongoing policies. For example, the transfer of dopamine responses from unconditioned to conditioned stimuli reflect the increase in confidence about “*what I should do*” after observing a conditioned stimulus. In short, the reinforcement learning (reward learning) story associates dopaminergic responses with RPE, while the active inference story treats dopaminergic function as encoding the confidence in policy selection, based upon inferred states of the world.

### Homeostatic reinforcement interface

Despite the paucity of direct evidence for the interface between homeostatic variables and reward or precision computations, there is convergent (but still tentative) evidence to suggest how the interface could be implemented (Fig. 6a & 6b). VTA-DA neurons host a number of receptors that would mediate this interface; they are positively modulated by ghrelin, a hormone reporting short-term energy deficits, and melanocyte-stimulating hormones (α,β,γ) released from POMC neurons; whereas they are inhibited by AgRP and its co-transmitter GABA, as well as by hormone insulin, and leptin, as well as by GLP-1 (Ferrario et al. 2016). Thus, the cells themselves provide ample opportunity for interfacing from homeostatic state information to the precision or reward value signal that is broadcast to the mesolimbic system from the VTA.

Thus, the first question to ask is do they connect directly? Yes. AgRP axons project directly to both the VTA and the substantia nigra (Dietrich et al. 2012), and POMC neurons have been labelled by retrograde tracers in the VTA (King & Hentges 2011, Fig. 7). On the reinforcement learning account AgRP, neurons should positively modulate VTA-DA, since they encode something hunger-like and hunger increases the value of food. Conversely, POMC neurons, encoding the converse of AgRP, should then negatively modulate VTA-DA.

In fact, the opposite appears to be observed. As noted above, AgRP neurons exert inhibitory effects over VTA-DA cells, directly via inverse agonism of the MCR3 receptor (the predominant melanocortin receptor expressed on VTA-DA neurons), and indirectly via its co-transmitter GABA, that acts to stimulate inhibitory interneurons that inhibit VTA-DA cells. Symmetrically, POMC neurons release melanocyte-stimulating hormones which also activate MCR3, which activates the VTA-DA neurons. These empirical results fit comfortably with active inference in the following sense: If AgRP neurons encode the hypothesis that “*I need to eat*”, then higher level (allostatic) expectations about eating will suppress their activity. However, the higher-level expectations that “*I am about to eat*” must be held with confidence or precision that is accompanied by dopaminergic discharges. In short, when AgRP firing is suppressed this will necessarily entail a confidence belief that “*I am about to eat*” and a disinhibition of dopaminergic outflow to the cortical basal ganglia thalamic systems responsible for policy selection.

### Why the counterintuitive responding?

Taken in the context of the predictive control findings discussed above if AgRP is deactivated, this means that under our interpretation, the precision on the prediction of positive future energy wealth increases, which is encoded via phasic dopamine, via the release of VTA-DA from inhibition. Likewise, if the POMC neurons are simultaneously activated by the same sensory evidence, then this has an excitatory effect on the same VTA-DA cells, which together with the AgRP disinhibition, provides a means by which VTA-DA signalling can be anchored to updates of predictions on future energetic wealth (i.e., the consequences of beliefs about the current long-term policy are assigned high precision or confidence). This is corroborated by ex-vivo recordings in which VTA-DA neurons increase baseline firing to injections of γ-MSH (Pandit et al. 2015). It should be noted that these findings seem to be at odds with the existing consensus that AgRP neurons acts to increase feeding and reward, and MSH acts to decrease feeding (Yen & Roseberry 2015).

This might however be an artefact of the way these injection experiments are performed. Pandit and colleagues (2015) show that infusion of a non-specific MCR agonist that targets both MC3R and MC4R, then sucrose responding decreases (also shown by Shanmugarajah et al. 2017; Yen & Roseberry 2015). However, by adding an MC4R antagonist, turns this response into an increase in sucrose responding. The important point here being that MC3R is predominantly expressed in the VTA, whereas the MC4R is expressed more broadly (for instance in the Nucleus accumbens) but not in the VTA. Interestingly the MC3R are predominantly expressed on the D2R expressing neurons that project into the nucleus accumbens. Notably the increased sucrose responding mediated by MC3R receptor agonism, is dependent on dopamine since DA-antagonism eliminates the effect (Pandit et al. 2015). Together, this might explain the apparent contradictions between prior work showing that melanocortin injections decrease responses to food reward (Yen & Roseberry 2015).

## Conclusion

In addressing the problems of homeostatic control, we have tried to bridge between several different fields, from evolutionary theory, neo-behaviourism, reinforcement learning, and computational and metabolic neurosciences. In doing so, this paper offers two main contributions.

First, we revisited how control theory and reinforcement learning have been applied to motivational behaviour, reward and homeostasis. We marshalled existing as well as novel arguments, for how these schemes rest on biologically implausible assumptions, either via circular definitions of reward, unprincipled groundings of value or drive functions, or by assuming degrees of certainty that are incompatible with the capricious nature of our natural habitats. Against this background, we have reviewed the active inference framework as it applies to these same homeostatic control problems. Putatively, we conclude that this offers promise in circumventing the shortcomings summarised above, and at the same time retains and builds on several important notions from comparator-based and reinforcement learning models. Of these conceptual advances, the most important are that set points are absorbed into prior beliefs about hidden viscero-sensory states, that homeostatic errors are cast as precision-weighted errors on interoceptive predictions, and that optimal choice behaviour is framed as an inferential process given a generative model of the body and its environment.

Second, we reviewed extant evidence pertaining to the how homeostasis interfaces with value computations in the domain of nutrient energy. Focusing on the case of the arcuate nucleus, we reviewed recent evidence for its role in the predictive control of energy homeostasis, contextualising the observations in the context of competing theoretical formulations. We assembled evidence suggesting that a subset of these arcuate subpopulations project directly to, and are thus in a privileged position to opponently modulate, dopaminergic VTA cells as a function of energetic predictions over a spectrum of temporal horizons. Further, we have surveyed how circulating factors that contribute to the dynamics of glucose homeostasis are direct modulators of dopaminergic neurons in the midbrain as well. The emerging picture points to a multi-faceted homeostatic-reward interface between the hypothalamus and midbrain. This interface may play a pivotal role in the conjoint optimisation of physiological and behavioural homeostasis.

That said – given the current state of knowledge – assigning computational roles to hypothalamic neurons may be premature. The computational quantities entailed by active inference are many, and their differences can be subtle. For instance, whether AgRP or POMC are encoding predictions, prediction errors, interoceptive states, or precisions portended by those states, will require careful experimentation. Existing evidence does not yet conclusively support one or the other. However, we hope that the theoretical perspective offered here motivates empirical experiments that can disambiguate between computational formulations of the brain’s homeostatic-reinforcement interface.

## Acknowledgements

We thank Mehdi Keramati and Boris Gutkin for several helpful discussions. This work was supported by the following funders: H.R.S (Lundbeck Foundation Grant of Excellence “ContAct” ref: R59 A5399; Novo Nordisk Foundation Interdisciplinary Synergy Programme Grant “BASICS” ref: NNF14OC0011413) O.J.H (Lundbeck Foundation, ref: R140–2013–13057; Danish Research Council ref: 12–126925) T.M. (Lundbeck Foundation ref: R140–2013–13057), K.F (The Wellcome Trust ref: 088130/Z/09/Z), D.B (The Francis Crick Institute, which receives its core funding from Cancer Research UK, the UK Medical Research Council, and the Wellcome Trust).

## Author contributions

O.J.H and T.M conceived of the paper and made the figures. All authors contributed to the writing and editing of the paper.

## Author Information

The authors declare no competing financial interests.

